# Exploring the *in-situ* evolution of Nitrofurantoin resistance in clinically derived Uropathogenic *Escherichia coli* isolates

**DOI:** 10.1101/2022.07.19.500598

**Authors:** Maxime Vallée, Chris Harding, Judith Hall, Phillip D Aldridge, Aaron Tan

**Affiliations:** Biosciences Institute, Faculty of Medical Sciences, Newcastle University, UK; Department of Urology, Poitiers University Hospital, Poitiers, France; Translational and Clinical Research Institute, Faculty of Medical Sciences, Newcastle University, UK; Urology Department, Freeman Hospital, Newcastle upon Tyne Hospitals NHS Foundation Trust, UK

**Author notes:** Co-Corresponding authors: P.D.A.; A.T. SCELSE, Nanyang Technological University, SBS-01N-27, 60 Nanyang Drive, 637551, Singapore.

## Abstract

**BACKGROUND:** Nitrofurantoin has been re-introduced as a first-choice antibiotic to treat uncomplicated acute urinary tract infections in England and Wales. Its mode of action involves initial reduction by nitroreductases, to generate electrophilic intermediates that inhibit protein and nucleic acid synthesis. Highly effective against common uropathogens such as *Escherichia coli*, its use is accompanied by a low incidence (<10%) of antimicrobial resistance. Resistance to Nitrofurantoin is predominantly via the acquisition of loss-of-function, step-wise mutations in the nitroreductase genes *nfsA* and *nfsB*.

**OBJECTIVE:** To explore the *in situ* evolution of Nit^R^, longitudinal uropathogenic *E. coli* isolates recovered from two rUTI patients.

**RESULTS:** Growth rate analysis identified a 2-10% slower doubling time for Nitrofurantoin resistant strains, but statistically, these data suggested there was no fitness advantage of evolved strains over their sensitive predecessor (ANOVA P-value = 0.13). Genetic manipulation of *E. coli* to mimic nitrofurantoin resistance evolution, again confirmed no fitness advantages (ANOVA P-value = 0.22). Rather, further analysis argued that a first-step mutant gained a selective advantage, at sub-MIC (4-8 mg/L) nitrofurantoin concentrations.

**CONCLUSION:** Correlation of these findings to Nitrofurantoin pharmacokinetic data suggests that the low incidence of *E. coli* Nit^R^, within the community, is driven by urine-based nitrofurantoin concentrations that selectively inhibit the growth of *E. coli* strains carrying the key first-step loss-of-function mutation.

## Introduction

Nitrofurantoin is a broad-spectrum antibiotic that has been used clinically since the mid-1950s to manage uncomplicated urinary tract infections (UTIs)^1^. With the introduction of trimethoprim and modern β-lactam antibiotics, which are more suitable for individuals with decreased renal function, its popularity waned in the 1970s ^1,2^. However, due to increased trimethoprim and β-lactam resistance – with trimethoprim resistance being reported in 30-35% of *Escherichia coli* isolates associated with UTI - the 2015 England and Wales NICE recommendations replaced trimethoprim with nitrofurantoin as the front-line antibiotic treatment for uncomplicated lower UTIs ^3^. Reassuringly, increases in nitrofurantoin prescriptions have not only been associated with decreased trimethoprim resistance, but no apparent changes in nitrofurantoin resistance. Public Health data supports nitrofurantoin resistance remaining, consistently, below 10% ^4^.

Nitrofurantoin is effective against a variety of common uropathogenic bacterial species, including *E. coli, Enterococcus spp*., *Enterobacter spp*. and *Klebsiella spp*. ^5,6^. However, the exact mechanism(s) by which nitrofurantoin exerts its antimicrobial effects is still under investigation. It has been established that many uropathogens carry genes encoding oxygen insensitive nitroreductases that convert nitrofurantoin into electrophilic intermediates that attack bacterial ribosomal proteins, thereby inhibiting protein synthesis and facilitating microbial death ^5^. When nitrofurantoin is present at high concentrations these intermediates can also interfere in nucleic acid synthesis and aerobic metabolism targeting the citric acid cycle ^5^. This killing mechanism is supported by the fact that uropathogens e.g. *Proteus mirabilis*, that naturally lack the nitroreductase genes are resistant to nitrofurantoin ^5,7^.

The low incidence of nitrofurantoin resistance was originally attributed to the ability of the drug, once metabolised, to interfere with multiple metabolic mechanisms ^8^. However, bacterial nitroreductases are not essential proteins, which allow bacteria to acquire chromosomally derived loss-of-function mutations generating the nitrofurantoin resistance (Nit^R^) phenotype ^9^. In *E. coli*, for example, the inactivation, via deletion or point mutation, of the chromosomally located genes, *nfsA* and *nfsB*, which account for 70% and 30 % of nitroreductase activity respectively, appear a common mechanism of gaining Nit^R 9^. Other pathways leading to Nit^R^ include the horizonal acquisition of the efflux system *oqxAB* or chromosomal mutations in *ribE* ^10^.

The inactivation of *nfsA* and *nfsB* follows a two-step evolutionary pathway: *nfsA* before *nfsB* ^11^. Supporting this evolutionary pathway, genetic surveillance studies of Nit^R^ uncovered *nfsA*^*-*^ *nfsB*^*+*^ isolates, but seldom the reciprocal *nfsA*^*+*^ *nfsB*^*-*^ combination ^10,12^. *In vitro* experiments have also demonstrated that Δ*nfsA* single mutants exhibit intermediate levels of nitrofurantoin resistance (MIC < 64 mg/L) compared to their parental isolates ^12^. Yet, despite its increased use, *E. coli* resistance to nitrofurantoin remains consistently below 10% ^4^. Sandegren et al (2008) provided insight into potential factors contributing to the low incidence of Nit^R^ among *E. coli* isolates. These authors reported the growth rate of clinical Nit^R^ isolates, characterised by mutations in both *nfsA* and *nfsB*, to be 6% slower compared to clinical Nit^S^ isolates ^8^. These data were supported by *in vitro* studies where spontaneous Nit^R^ mutants showed a 1-3% slower growth rate compared to their parental strain. Sandegren et al (2008) thereby concluded that despite the nitroreductase being presumed as non-essential, there was a fitness cost to uro-associated *E. coli* of inactivating *nfsA/nfsB* ^8^. The authors argued that loss of fitness impacted the establishment of resistant isolates in the urinary tract, thereby explaining the low incidence of nitrofurantoin resistance among urinary *E. coli* isolates ^8^. Additionally, all their resistant isolates were unable to grow at 200 mg/L nitrofurantoin, proposed by the authors to be the minimum nitrofurantoin concentration in the urinary tract during treatment ^8^. A recent pharmacokinetic study, has however, reported the maximum urinary nitrofurantoin concentration during treatment to be in the region of 94 mg/L ^13^, two-fold lower than the presumed minimum urinary nitrofurantoin concentration used by Sandegren et al. (2008), which helps explain the isolation and persistence of Nit^R^ isolates clinically.

While genome surveillance studies support Nit^R^ being linked to point mutations and deletions of the *nfsA/nfsB* genes ^10,12^, they are unable to explain the factors that drive resistance phenotype selection. Understanding these factors are key to informing prescribing guidelines that underpin UTI treatments and future urology antibiotic stewardship programmes. One potential approach to investigate such mechanisms is to exploit longitudinal uro-associated isolates recovered from individual rUTI patients presenting with Nit^R^ who have been treated either acutely or prophylactically with nitrofurantoin. AnTIC was an open label randomised controlled trial that assessed the efficacy of antibiotic prophylaxis in reducing the incidence of symptomatic UTIs in clean intermittent self-catheterised patients over a period of 12 months ^14^. During the trial, uro-associated *E. coli* isolates from individual patients suffering rUTIs were banked. Forty-five isolates from 17 patients experiencing persistent *E. coli* colonisation were analysed by whole-genome sequencing ^15^ and five isolates from two patients (PAT1646 and PAT2015) supported Nit^R^ acquisition during the trial period. Clinical data associated with these cases confirmed exposure to nitrofurantoin. Of these two cases, one provided an opportunity to investigate the *in situ* evolutionary dynamics driving nitrofurantoin resistance and address aspects of the Nit^R^ phenotype that underpin the low incidence of Nit^R^ observed among community isolates of *E. coli*.

## Materials and Methods

### Bacterial strain and growth conditions

Strains used or constructed in this study are shown in supplementary materials **(Table S1)**. Overnight cultures were grown in LB (recombination/transformation) or MHB (MIC/growth/viability assays) media at 37°C with constant shaking at 160 rpm. Antibiotics for selection were used at concentrations that have been previously described ^16^.

### pBKK plasmid construction

Five pBKK plasmids carrying different DNA constructs of the *nfsA* or *nfsB* region (*nfsA* 1646B, Δ*nfsA*, Δ*nfsA-rimK*, Δ*rimK* and Δ*nfsB*) were generated using Gibson assembly, with pBKK as the vector backbone. The pBKK vector was constructed using Gibson assembly, with pBlueScript II SK (Ampicillin resistant) as the vector backbone and the kanamycin resistance gene from pKD4 as the insert. Since 1646 BASE isolates are innately ampicillin resistant, pBKK plasmids enabled the selection of transformed 1646 BASE isolates. All plasmids were sequenced, and primers are defined in **Table S2**.

### Mutant generation

*nfsA/nfsB* mutants were generated using a two-step recombination strategy involving the lambda-red and CRISPR systems ^17,18^. Lambda-red mediated recombination was first used to replace the *nfsA/nfsB* region with *cat* from pWRG100 ^19^, encoding chloramphenicol resistance. The *cat* gene was both a selective marker and a unique genomic target for the CRISPR sgRNA ^20^. During CRISPR mutagenesis, *cat* was replaced with a DNA construct carrying a modified version of the *nfsA/nfsB* region (Δ*nfsA*, Δ*nfsA-rimK*, Δ*rimK, nfsA* T37M, Δ*nfsB*, Δ*nfsB30*). Where necessary the target mutant was amplified from the pBKK plasmids or directly from chromosomal DNA (*nfsA* T37M and Δ*nfsB30*). Double mutants were generated by first replacing the *nfsA* region with the appropriate DNA construct followed by the *nfsB* region. Lambda-red expression on the pCas9 plasmid was induced with 0.1% arabinose when cultures reached an OD_600_ = 0.1. Cells were prepared for electroporation once the induced cultures had reached an OD_600_ = 0.6–0.8. Colonies were checked for insertion of the appropriate DNA constructs through colony PCR and sequencing.

### MIC assay

MIC assays were performed using sterile Greiner Bio-One clear 96-well flat-bottom plates. Bacterial isolates were grown overnight in MHB. Subcultures were generated by diluting overnight cultures 1 × 10^−4^ with MHB. For each isolate, 100 µl of subculture was pipetted into every well of a single row of the plate. The final nitrofurantoin concentration in each row ranged from 0-128 mg/L. The plates were covered with Breathe-Easy sealing membrane following manufacturer’s instructions before being incubated statically for 16 hours at 37°C in a SANYO incubator. After incubation, the membrane was removed, and bacterial growth determined using a BMG Labtech FLUOstar OPTIMA microplate reader set up for OD_600_ measurement.

### Growth assay and analysis

Growth curve assays were performed using sterile Greiner Bio-One clear 96-well flat-bottom plates. Four independent colonies from each isolate to be tested were grown overnight each in 3 ml of MHB. The OD_600_ of 10 random cultures was measured and averaged to generate the mean OD_600_ value for the entire batch of isolates. This mean OD_600_ value was used to determine the volume of overnight culture required to generate 2 ml subcultures (MHB) with a starting OD_600_ = 0.02 for each isolate/colony. A 200 µl volume of each subculture was pipetted into each well of a 96-well plate. Three wells acted as controls and each contained only MHB. The plate was covered with Breathe-Easy sealing membrane before being inserted in a BMG Labtech FLUOstar OPTIMA microplate reader. The plate was incubated for a period of 10.2 hours (90 timepoints) at 37°C, orbital shaking = 250 rpm, with OD_600_ readings taken every 400 seconds.

The maximum growth rate of each isolate was determined by calculating the maximum slope (gradient). OD_600_ readings used for slope calculations were converted to natural log. The slope was calculated using a moving “frame” of 8 timepoints ^21^. The outcome was a data frame of slope calculations from which the maximum slope could be determined. The doubling time of each isolate was calculated by dividing the natural log of 2 by the maximum slope. Statistical analysis of the doubling times for all growth experiments was performed using either T-Tests or ANOVA. When T-Tests were conducted the Bonferroni correction was considered to determine the threshold P-value for significance.

### Viability assay

Overnight bacterial cultures were prepared and following OD_600_ measurements were diluted to achieve a starting density of 50-100 CFU/ml in 6 ml of MHB media and incubated for 16 hours at 37°C. One hundred microlitres of serial dilutions of the overnight cultures (10^−5^ and 10^−6^) were plated onto a CPSE plates. CPSE plates were incubated overnight at 37°C, photographed and colonies counted with the image processing package ImageJ.

### Ethical approval

Permission to use clinical isolates and data from the AnTIC clinical study was derived from prior ethical approvals (Ethics: 19/NS/0024, IRAS Project ID: 243903, Ref: 2586/2016) ^15^.

## Results

### *In situ* development of nitrofurantoin resistance amongst clinical *E. coli* isolates

Within AnTIC, 45 uro-associated isolates from 17 patients with persistent *E. coli* colonisation were previously analysed using whole-genome sequencing (WGS) ^15^. Five isolates from two patients (PAT1646 and PAT2015) provided examples that supported *in situ E. coli* acquisition of nitrofurantoin resistance following exposure to the antibiotic. MLST and cgMLST genotyping confirmed stable *E. coli* colonisation in both patients during the participation in AnTIC (**Fig. 1 A**). PAT1646 was colonised by ST131 whereas PAT2015 was colonised by ST58 (**Fig. S1**). These *E. coli* isolates, developed nitrofurantoin resistance by acquiring inactivating mutations in *nfsA* and *nfsB* respectively (**Fig. 1 B**). The Nit^S^ (MIC: 8 mg/L) baseline isolate of PAT1646, 1646 BASE, possessed wild-type *nfsA* and *nfsB* genes. The 6-month isolate, 1646 6, was Nit^R^ (MIC 128 mg/L) and had a T37M point mutation in *nfsA* and a complete deletion of *nfsB*, spanning 30 Kb upstream of the start codon. The baseline isolate of PAT2015, 2015 BASE, was Nit^S^ (MIC: 32 mg/L). It was characterised by a complete deletion of *nfsA*, but an intact, wild-type *nfsB* (**Fig. 1 B**). The 12-month isolate, 2015 12, was Nit^R^ (MIC: 128 mg/L) and characterised by the complete deletion of *nfsA* and a partial deletion in *nfsB* (**Fig. 1 B**).

**Figure 1.**
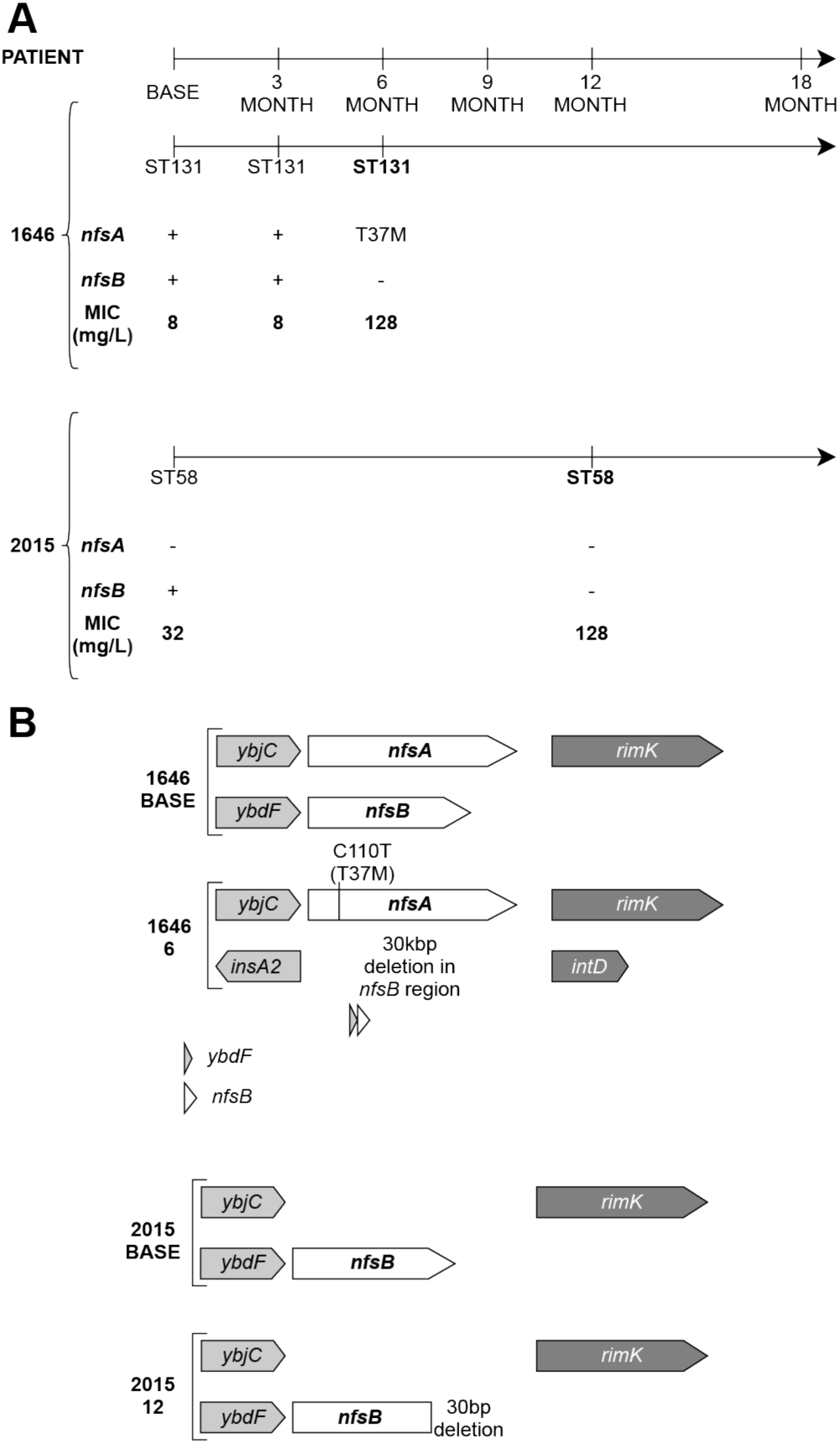
Acquisition of the Nit^R^ phenotype in PAT1646 and PAT2015. **A:** Timeline for each patient indicating the temporal isolation of each strain, the sequence type identified via MLST genotyping and the MIC for each isolate. **B:** *nfsA* and *nfsB* genotype for the BASE and Nit^R^ isolates from PAT1646 and PAT2015.

Growth kinetics of the BASE strains, 1646 6 and 2015 12 isolates resulted in comparable growth curves (**Fig 2 A**) which suggested no difference in bacterial fitness. Calculation of the doubling time from the growth rates defined comparable times for all four strains, ranging between 20.8 ± 0.7 mins for 1646 BASE and 23 ± 0.8 mins for 1646 6 (ANOVA P-value = 0.13) (**Fig. 2 B**). Furthermore, in the absence of antibiotic, competition between 1646 BASE versus 1646 6 was minimal, reflected by a calculated competitive index of 0.22. These data therefore argued against a fitness advantage playing a significant role in driving the acquisition of Nit^R^ in rUTI patients treated with nitrofurantoin.

**Figure 2.**
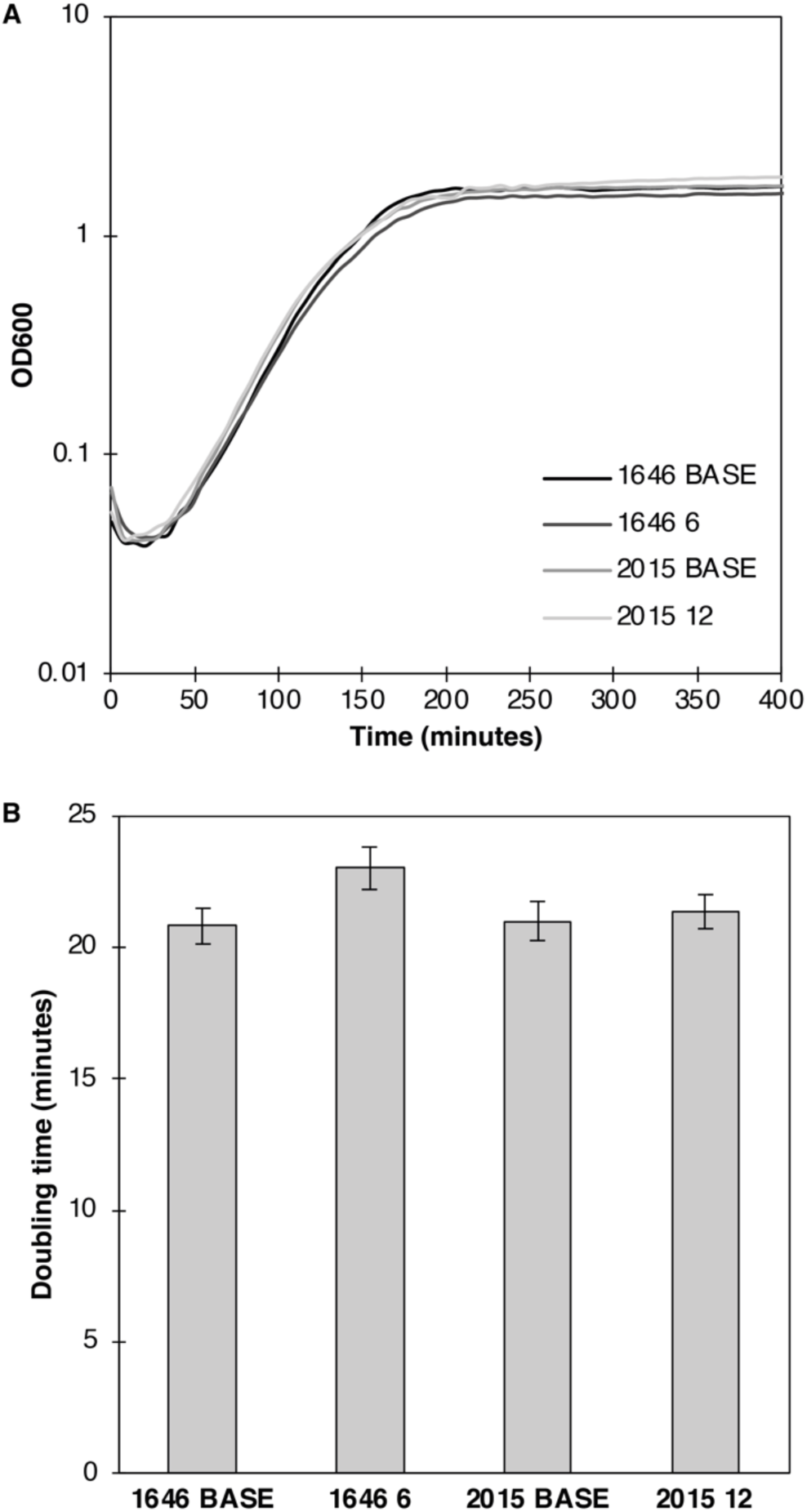
Growth characteristics of the BASE and Nit^R^ isolates from PAT1646 and PAT2015. **A:** Average growth curve data **B**. Maximal doubling time in minutes calculated from (**A**). The differences between the doubling times are consistent with previous studies of Nit^R^ but are not statistically significant (ANOVA P = 0.22). All data was calculated from n = 24 independent repeats of growth of each strain.

### Generating *nfsA* and *nfsB* mutants from Nit^S^ isolates

PAT1646 *E. coli* isolates 1646 BASE and 1646 6 represent a chronological “snapshot” of the same strain before and after developing nitrofurantoin resistance (**Fig. 1**). To investigate the underlying advantage / disadvantage of acquiring the Nit^R^ phenotype, mutants in *nfsA* and/or *nfsB*, were generated in 1646 BASE and the Nit^S^ laboratory *E. coli* strain W3110, a commonly used “wild-type” model strain ^22^. The *nfsA* and *nfsB* wild-type genes were inactivated via complete deletion (Δ*nfsA* and Δ*nfsB*) (**Fig. S2**). Using two isolates of contrasting genetic backgrounds (clinical vs laboratory), provided the opportunity to identify phenotypic effects, if any, of Δ*nfsA/ΔnfsB* combinations that were dependent or independent of genetic background. The *nfsA* and *nfsB* genes were inactivated via complete deletion (Δ*nfsA* and Δ*nfsB*) (**Fig. S2**). Mutants, Δ*nfsA* and Δ*nfsB*, were generated by targeted mutagenesis rather than *in vitro* selection to minimise the acquisition of spontaneous, compensatory mutations linked to nitrofurantoin exposure, which could distort the phenotypic behaviour of Δ*nfsA* / Δ*nfsB* mutations.

The *rimK* gene is in an operon with *nfsA* (**Fig. 1B**) and encodes an enzyme involved in the post-translational modification of ribosomal protein S6 via the addition of glutamate residues ^23^. There is evidence of clinical Nit^R^ isolates that carry partial deletions in *rimK*, therefore Δ*rimK* and Δ*nfsA-rimK* mutations were also created (**Fig. S2**) ^15^. Using the flexibility of CRISPR/Cas technology to generate the mutants also allowed 1646 BASE to be engineered to model the impacts of the 1646 6 *nfsA* mutation (*nfsA* T37M) and the 30 Kb *nfsB* deletion (Δ*nfsB30*). However, only *nfsA* T37M was modelled in W3110 (**Fig. S2**).

### Characterisation of Δ*nfsA / ΔnfsB* mutants

The Nit^R^ phenotypes of the strains generated were quantitatively assessed via MIC assays and data shown in **Fig. 3**. Isolates with a MIC ≥ 64 mg/L were classified as Nit^R 24,25^ and strains 1646 BASE, W3110 and 1646 6 were included as Nit^S^ and Nit^R^ controls. Single 1646B mutants with Δ*nfsA* or *nfsA* T37M supported MIC values of 16 mg/L thus still clinically Nit^S^ (**Fig. 3**). Single mutants Δ*rimK* and Δ*nfsB* single mutants exhibited a MIC comparable to their parental background. A similar pattern was observed for the W3110 although the Nit^S^ MIC of the parent and *nfsA*^+^ variants was 4 mg/L compared to 8 mg/L in 1646 BASE (**Fig. 3**).

**Figure 3.**
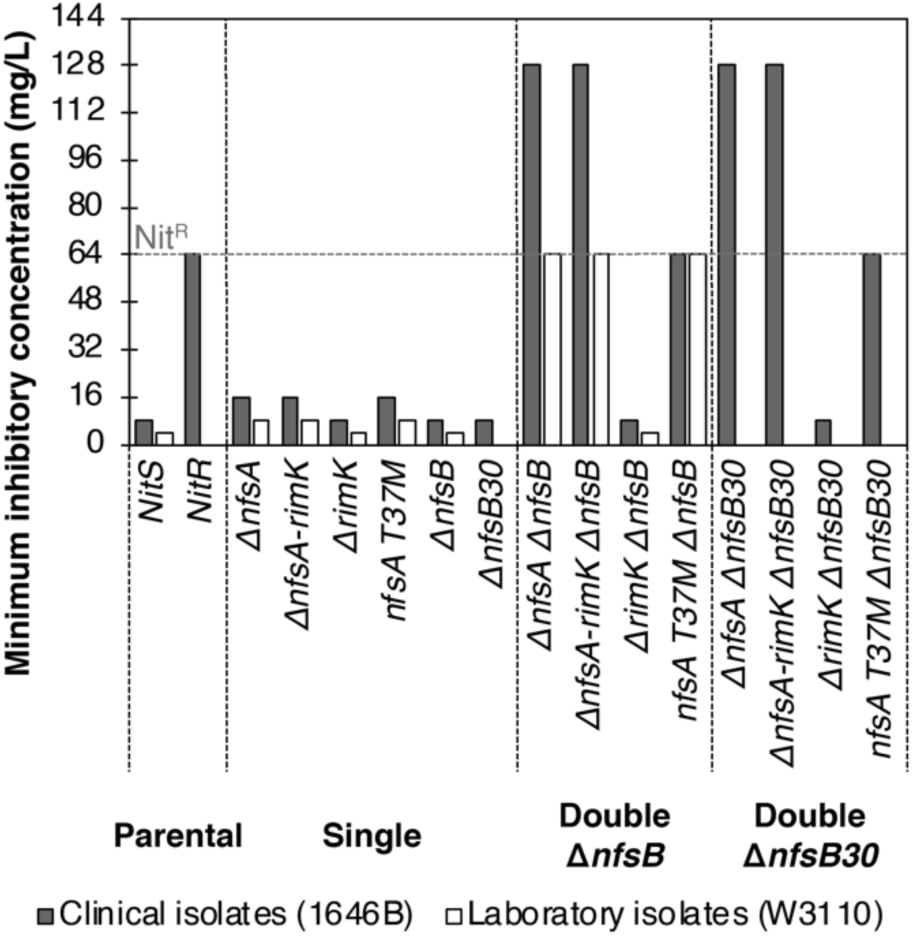
Minimum Inhibitory Concentrations (MIC) of Δ*nfsA* and Δ*nfsB* mutant combinations in 1646 BASE (grey bars) and the laboratory strain W3110 (white bars). The grey dashed horizontal line indicates the nitrofurantoin concentration defined as being the point at which strains are defined as clinically resistant to the antibiotic. Data represent a minimum of 3 independent repeats of the assay for each strain.

*In vitro* 1646 6 and its corresponding engineered mutant (*nfsA* T37M Δ*nfsB*) showed MICs of 64 mg/L respectively. In contrast, 1646 BASE double mutants in which *nfsA* was inactivated via complete deletion (Δ*nfsA* Δ*nfsB*, Δ*nfsA-rimK* Δ*nfsB*) displayed higher MICs (128 mg/L) (**Fig. 3**). The MICs of the *nfsA*^+^ Δ*rimK* Δ*nfsB* variants were comparable to their single and parental derivatives (8 mg/L or 4 mg/L). Collectively, these data indicated that only single mutants with inactivated *nfsA* had a modest 2-fold MIC increase when compared to their parental isolate while double mutants, including Δ*nfsA*, had substantially higher MICs.

### Plasmid-based complementation of Nit^R^ Δ*nfsA* Δ*nfsB* mutants

To verify that inactivating *nfsA* was critical in the acquisition of Nit^R^, the engineered 1646 BASE and W3110 Δ*nfsA* Δ*nfsB* mutants were transformed with modified pBlueScript II SK (pBKK) plasmids carrying genetic variants of the *nfsA* region (**Fig. 4 A**). The pBKK empty vector was used as a negative control. MIC assays were performed on transformed strains (**Fig. 4 B**). Complementation was only observed when pBKK plasmids carried functional *nfsA*. Consistent with the role that *nfsA* plays in the Nit^R^ phenotype, complementation led to a strong 32-fold reduction in MIC compared to the negative control (**Fig. 4 B**).

**Figure 4.**
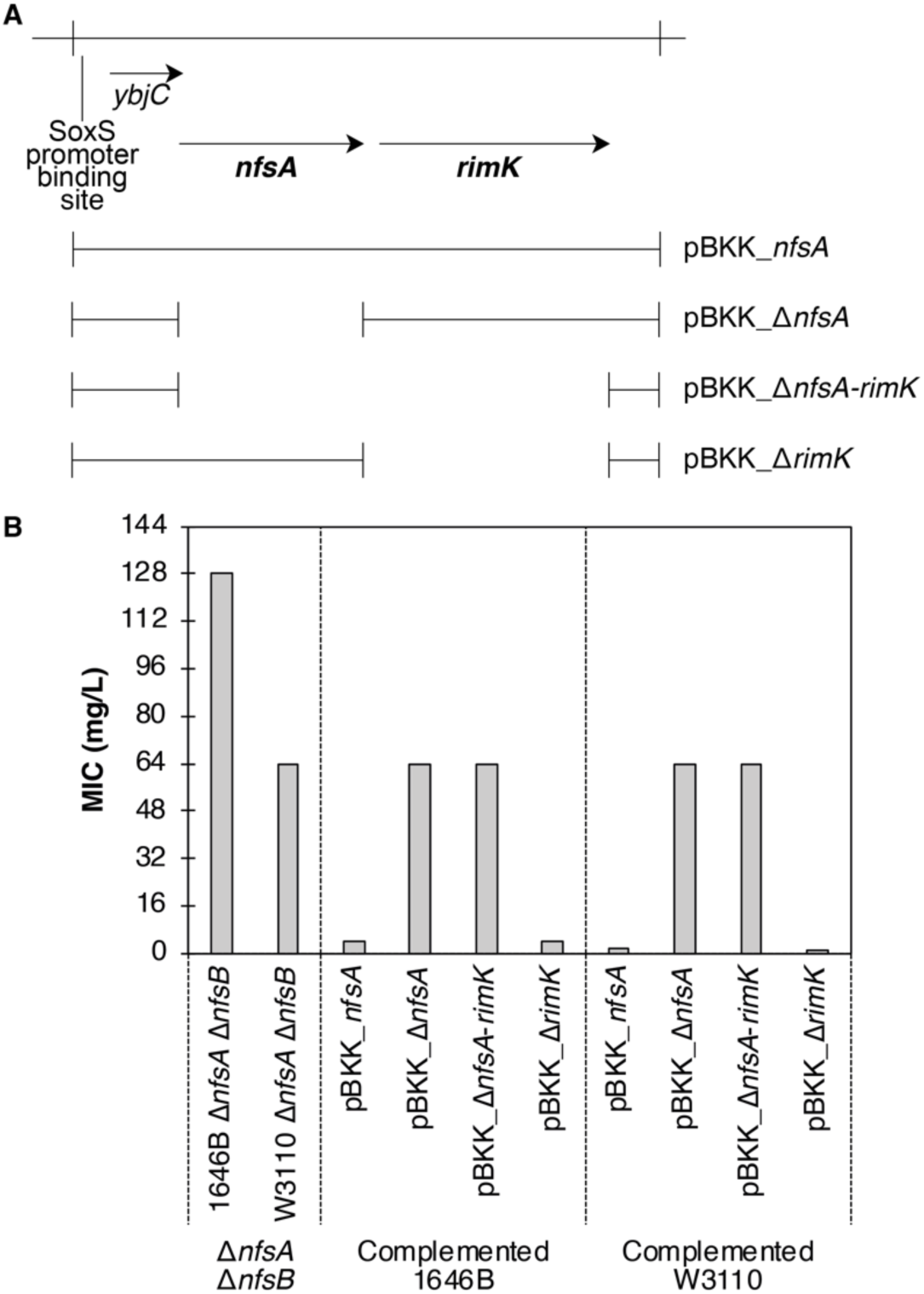
Complementation of Δ*nfsA* with ectopic expression of *nfsA* from pBKK. **A:** Schematic diagram of plasmid constructs used in these complementation assays. **B:** MIC data for strains used. Data represent a minimum of 3 independent repeats of the assay for each strain.

### Fitness of *nfsA/nfsB* mutants

The relative fitness of the mutants was determined by maximum bacterial growth rate in terms of doubling time, with an assumed increase in fitness corresponding to shorter doubling times. In the absence of nitrofurantoin (0 mg/L), there was no significant difference in doubling time between any of the mutants and their corresponding parental isolates (ANOVA 1646 BASE: P-value = 0.22; W3110: P-value = 0.46) (**Tables 1 and 2**). Pairwise T-Test analysis using a Bonferroni corrected significance threshold of P = 0.0009 agreed with the ANOVA analysis (**Tables S3 and S4**).

**Table 1.** Doubling times of mutants and clinical isolates relating to PAT1646 with or without nitrofurantoin

| Strains | Nitrofurantoin concentration (mg/L)* |  |  |  |  |  |  |  |
| --- | --- | --- | --- | --- | --- | --- | --- | --- |
|  | 0 | 2 | 4 | 8 | 16 | 32 | 64 | 128 |
| 1646 BASE | 21.9 ± 0.9 | 24.7 ± 0.7 | 29.3 ± 1.1 | 43.2 ± 1.8 |  |  |  |  |
| 1646 6 | 25.7 ± 1.1 | 25.1 ± 0.6 | 24.5 ± 1.2 | 25.3 ± 1.1 | 27.9 ± 0.9 | 35.2 ± 1 | 142 ± 11.3 |  |
| 1646B $\Delta nfsA$ | 24.2 ± 0.6 | 24.3 ± 0.9 | 23.6 ± 1 | 31 ± 2 | 60.1 ± 1.6 | | | |
| 1646B $\Delta nfsA$-*rimK* | 23.7 ± 0.9 | 26.2 ± 1 | 25.4 ± 1.3 | 32.2 ± 1.9 | 64.7 ± 2.4 | | | |
| 1646B *nfsA* T37M | 22.5 ± 0.8 | 24.2 ± 0.9 | 24.1 ± 1 | 30.3 ± 2.1 | 55.4 ± 2.2 |  |  |  |
| 1646B $\Delta rimK$ | 24.1 ± 1.2 | 25.3 ± 0.4 | 28.3 ± 1.3 | 45.9 ± 1.4 | | | | |
| 1646B $\Delta nfsB$ | 22.2 ± 1.1 | 25.9 ± 0.8 | 29.6 ± 1 | 45 ± 0.9 | | | | |
| 1646B $\Delta rimK$ $\Delta nfsB$ | 21.8 ± 1.8 | 25.3 ± 1.2 | 30.5 ± 0.8 | 43.7 ± 1.6 | | | | |
| 1646B $\Delta nfsA$ $\Delta nfsB$ | 25.7 ± 0.4 | 25.6 ± 0.9 | 25 ± 0.5 | 24.3 ± 1 | 26.6 ± 0.3 | 34.2 ± 1.3 | 90.5 ± 5.9 | |
| 1646B $\Delta nfsA$-*rimK* $\Delta nfsB$ | 21.9 ± 1.6 | 23.4 ± 0.8 | 22.6 ± 0.7 | 23.4 ± 1.3 | 27.7 ± 0.9 | 33.7 ± 0.8 | 88 ± 3.4 | |
| 1646B *nfsA* T37M $\Delta nfsB$ | 24.2 ± 0.8 | 25 ± 1.1 | 24.7 ± 0.5 | 27.7 ± 1 | 28.7 ± 0.5 | 37.6 ± 0.9 | 131.6 ± 10.8 | |

\* Grey zones identify significant changes in the doubling time (**Table S3**)

**Table 2.** Doubling times of mutants and the parental strain W3110 with or without nitrofurantoin

| Strains | Nitrofurantoin concentration (mg/L)* |  |  |  |  |  |  |  |
| --- | --- | --- | --- | --- | --- | --- | --- | --- |
|  | 0 | 2 | 4 | 8 | 16 | 32 | 64 | 128 |
| W3110 | 34.9 ± 0.8 | 37.8 ± 1 | 37.7 ± 1.2 | 74.7 ± 2.9 |  |  |  |  |
| 1646B $\Delta nfsA$ | 32.7 ± 1.7 | 35.4 ± 1.2 | 37.5 ± 1.8 | 50.9 ± 2.6 | | | | |
| 1646B $\Delta nfsA$-rimK | 33.1 ± 1.3 | 35.3 ± 1.1 | 40.4 ± 1.6 | 50.4 ± 2.1 | | | | |
| 1646B *nfsA* T37M | 29 ± 2.3 | 35.8 ± 1.1 | 38.8 ± 1.3 | 51.9 ± 2.4 |  |  |  |  |
| 1646B $\Delta rimK$ | 33.8 ± 1.2 | 40.3 ± 1.9 | 44.4 ± 2 | 79.5 ± 5.6 | | | | |
| 1646B $\Delta nfsB$ | 31.5 ± 3.1 | 34.9 ± 1.7 | 41.5 ± 1.4 | 73.7 ± 3.2 | | | | |
| 1646B $\Delta rimK \Delta nfsB$ | 35.8 ± 2.1 | 37.3 ± 1.4 | 39.9 ± 2.1 | 72 ± 4.3 | | | | |
| 1646B $\Delta nfsA \Delta nfsB$ | 31 ± 1.8 | 33.9 ± 0.8 | 36 ± 2.1 | 44.6 ± 1.6 | 39 ± 2.7 | 54.4 ± 1.9 | 117.2 ± 5.8 | |
| 1646B $\Delta nfsA$-rimK $\Delta nfsB$ | 32.5 ± 1 | 33.4 ± 0.9 | 34 ± 2 | 37.4 ± 1.6 | 32.8 ± 2.1 | 50.8 ± 1.4 | 97 ± 7.7 | |
| 1646B *nfsA* T37M $\Delta nfsB$ | 34.4 ± 0.5 | 37.7 ± 1.3 | 38.2 ± 2 | 44.1 ± 0.8 | 40.3 ± 1.3 | 56.7 ± 1.5 | | |

\* Grey zones identify significant changes in the doubling time (**Table S4**)

ANOVA analysis indicated that at 4 and 8 mg/L nitrofurantoin, there was significance within the bacterial growth data sets (P-value range < 0.0001 to 0.02) (**Tables 1 and 2**) with pairwise comparisons using T-Tests identifying specific trends. For example, at 4 mg/L nitrofurantoin an increase in doubling time from 21.9 ± 0.9 mins to 29.3± 1.1 mins was observed for the parental strain 1646 BASE with comparable increases for *nfsB* and *rimK* single mutants (**Table 1**). This increase became statistically significant across all comparisons at 8 mg/L where a doubling time of 43-46 minutes was observed (**Table 1: grey zones and Table S3**). The same trend was observed for W3110, which already had a lower intrinsic resistance to nitrofurantoin (**Table 2 and Table S4**). At concentrations >8 mg/L, the majority of these strains were unable to grow arguing that bacterial selection, rather than fitness, was the driving factor.

Similar data for the Δ*nfsA*, double mutants and the control strain 1646 6 was observed at higher concentrations of nitrofurantoin (**Table 1**). Interestingly, the higher the concentration of antibiotic the stronger the impact on growth with, for example, 64 mg/L leading to a 5-fold increase in doubling time for 1646 6. The 1646 BASE Δ*nfsA* Δ*nfsB* variant exhibited a 3.5-fold increase in doubling time at this concentration, consistent with the difference defined by the MIC analysis.

### Viability of *nfsA/nfsB* mutants

Growth experiments were initiated with an inoculation of between 0.01 – 0.05 OD600. This inoculum is equivalent to ∼ 1-5 x10^6^ CFU/ml, one log greater than the diagnostic threshold of microbes detected per milliliter of urine in an acute UTI (1 × 10^5^ CFU/ml) ^26^. It is well recognised that growth experiments can be influenced by the inoculum effect ^27^. Therefore, further growth analysis using a starting inoculum of 50 - 100 CFU/ml in 0 to 16 mg/L nitrofurantoin was investigated focussing on the viability of the 1646 BASE engineered strains rather than growth kinetics.

All strains tested grew well with or without 2 mg/L nitrofurantoin (**Table 3**). The 1646 BASE strain did not grow at 4 mg/L while the Δ*nfsB* mutant exhibited a significant (∼6 log-fold) reduction in viability (**Table 3: light grey zone**). Furthermore, single mutants of Δ*nfsA* and *nfsA* T37M behaved in a similar manner, but at double the antibiotic concentration i.e. 8 mg/L. All strains that possessed a Nit^R^ phenotype grew well at all nitrofurantoin concentrations tested (**Table 3**). These data strengthen the argument for selective advantage over fitness for step-wise *nfsA*^-^ intermediates (**Table 3**).

**Table 3.**
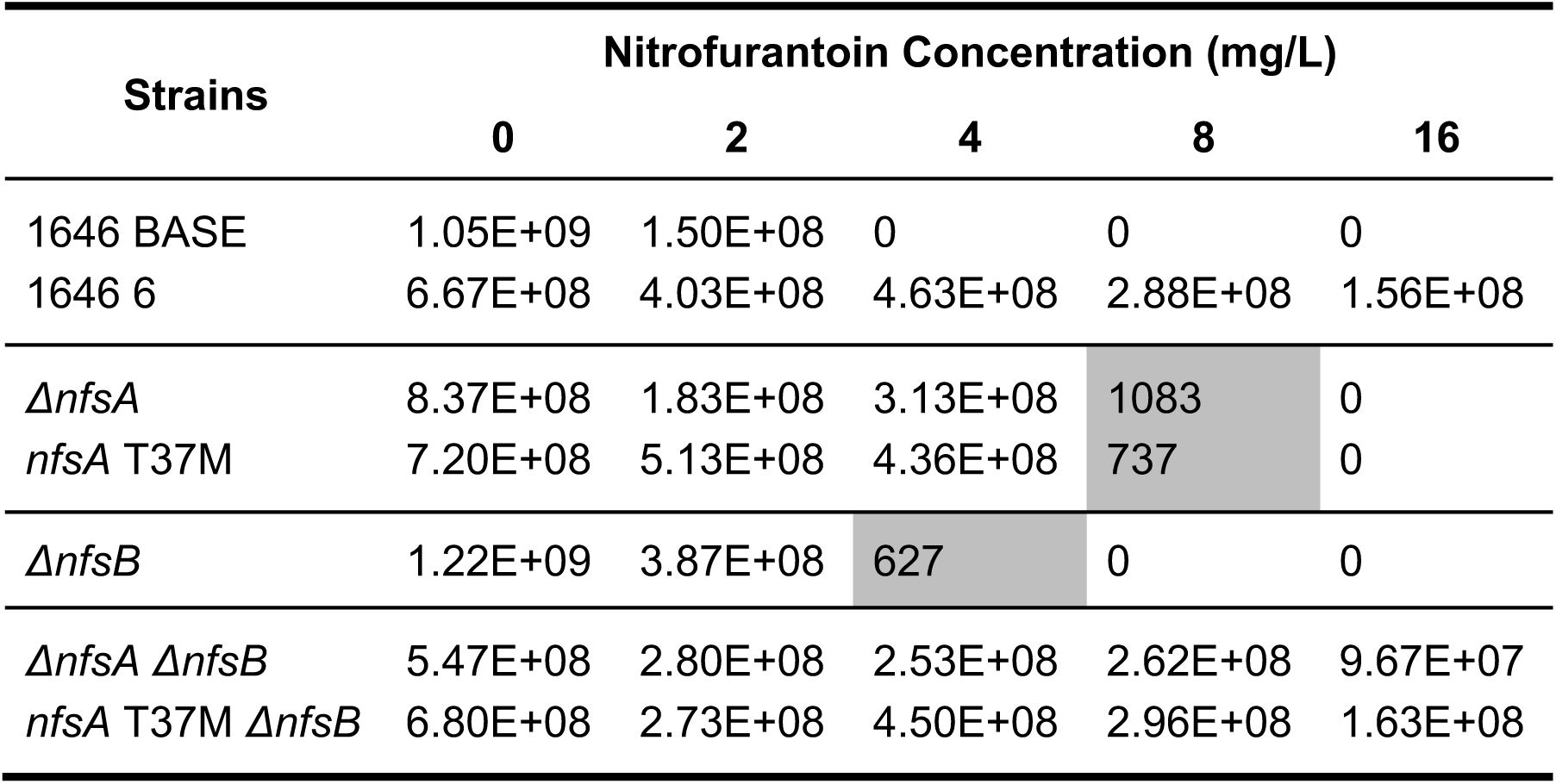
Viability of test strains in low inoculum growth experiments with sub-MIC concentrations of nitrofurantoin.

## Discussion

High level nitrofurantoin resistance requires the inactivation of two genes, which are located so far apart from each other (287kbp distance) that the likelihood of both being simultaneously inactivated through a single natural genetic event is non-existent^11^. Furthermore, there are multiple essential genes between *nfsA* and *nfsB*, which prevents them from being inactivated via a large single deletion. Hence inactivation of these genes is much more likely to occur in a stepwise manner, starting with *nfsA* followed by *nfsB*.

To explore the evolutionary mechanism(s) underpinning antibiotic resistance Nit^S^ and Nit^R^ uro-associated *E. coli* isolates recovered from rUTI patients treated with nitrofurantoin were exploited. These strains were unique as they reflected *in situ* evolution of *E. coli* over 6 to 12 months in a clinical environment from Nit^S^ *nfsAB*^+^ (PAT1646) or *nfsA*^-^*nfsB*^+^ (PAT2015) genotypes to resistant genotypes. Furthermore, these isolates are taken from an at-risk patient group that are treated regularly with antibiotics and thus are frequently exposed to antibiotic-induced selective pressure ^14,15^. Even so, the incidence of resistance evolution is low, suggesting a low risk in the general population. Consistent with the literature, the predominant mutations leading to Nit^R^ were deletions, although 1646 BASE to 1646 6 evolved Nit^R^ via the acquisition of a point mutation in *nfsA*: T37M. Growth and MIC data provided strong evidence that T37M, like other previously identified point mutations ^10,12^, was functioning as an inactivating mutation in *nfsA*. Data also suggested that the method by which *nfs* genes were inactivated (Δ*nfsA, nfsA*T37M, Δ*nfsB*, Δ*nfsB*30) did not significantly alter nitrofurantoin resistance. Essentially, gene inactivation *per se* was more important than the mode of inactivation. MIC data also suggested that *rimK*, that resides in an operon with *nfsA*, played no role in Nit^R^, at least in the 1646 patient isolates.

A previous study, Sandegren et al (2008), argued that fitness of Nit^R^ strains plays a key role in driving the low incidence of community AMR to nitrofurantoin. While the bacterial growth data reported in this study using strains generated from 1646 BASE and W3110 was consistent with Sandegren et al (2008), in that a small change in doubling times (2-10%) was observed, statistical analysis (ANOVA and pairwise T-Tests) argued that these growth changes were not significant and, alone, could not explain AMR. Bacterial resistance has traditionally been linked to the ability of isolates to grow in antibiotic concentrations that exceed the MIC of sensitive isolates. However, multiple studies have isolated resistant mutants from growth conditions where antibiotic concentrations were low enough for the proliferation of sensitive isolates ^28-30^. For most antibiotics, bacterial isolates experience some degree of growth reduction even at concentrations just below their MIC ^31-33^. Growth analysis data relating to 1646 BASE and W3110 strains and their genetic variants in the presence of Nitrofurantoin support this observation (**Tables 1 and 2**).

A sub-MIC selective window may be a key factor in driving antibiotic resistant phenotypes to successfully establish themselves within a population, and arguably, the wider the selective window the greater the incidence of resistance. However, these windows are not universal, with each being determined not only by the pharmacokinetics and pharmacodynamics of an antibiotic, but also the genetic background of a bacterial species / strain. The findings reported here suggest that the nitrofurantoin selective window in relation to rUTIs and uro-associated *E. coli* Nit^R^, is wide (> 8 mg/L), but this argument is only applicable when comparing Nit^R^ double mutants against their Nit^S^ parental isolates. From an evolutionarily perspective this comparison is biased as data suggests the emergence of Nit^R^ mutants is highly dependent on the ability of intermediate mutants with low level nitrofurantoin resistance to establish themselves within a population of sensitive isolates. In fact, comparing the growth of mutants (Δ*nfsA, nfsA* T37M) modelling these intermediate scenarios to their parental isolates (1646 BASE and W3110), argues for a very narrow selective window (4 to 8 mg/L). Despite these strong selective pressures favouring growth, bacteria may still fail to establish themselves in the urinary environment due to bladder voiding; bacterial survival necessitates a minimum doubling time of 36 minutes ^34^.

Clinically, urinary nitrofurantoin concentrations rarely fall to within this narrow concentration window of 4 to 8 mg/L, which limits the selection of these mutants ^13^. For example, extrapolating the urinary nitrofurantoin concentration data from Huttner et al. (2019), suggests the average urine concentration following a standard 50 mg dose may fluctuate between 20 - 40 mg/L during an 8-hour period ^13^. Essentially, these conditions inhibit intermediate mutants such as Δ*nfsA* and *nfsA* T37M establishing clinically, which in turn suppresses the emergence of double and hence Nit^R^ mutants. Therefore, the rarity of Nit^R^ intermediate mutants undermines the emergence of nitrofurantoin resistance in *E. coli*, explaining the low incidence of resistance.

## Acknowledgements

The authors would like to thank all participants of the original AnTIC trial who agreed to their samples being banked for further analyses.

## Transparency Declaration

None to Declare

---

**Table S1.**
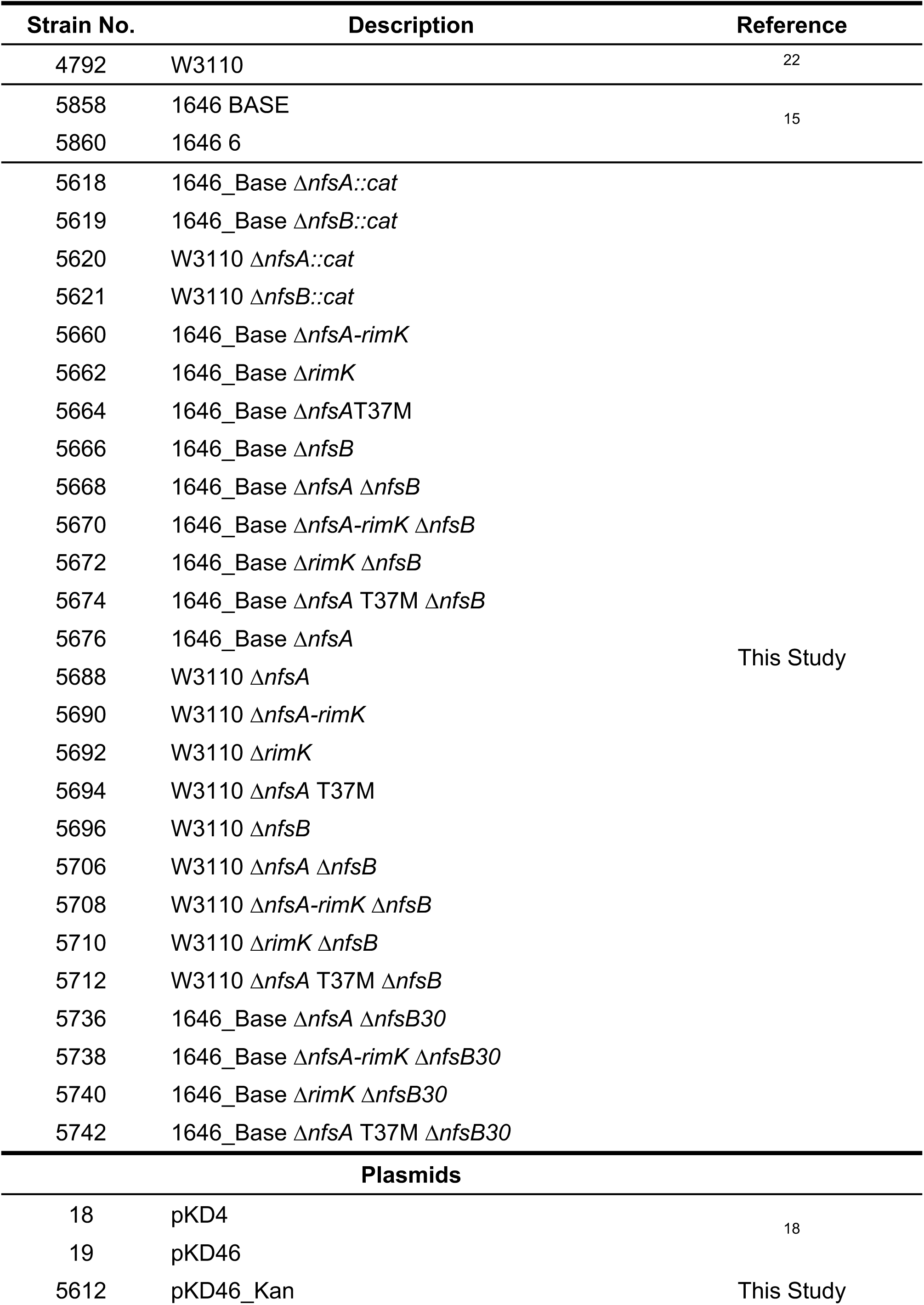
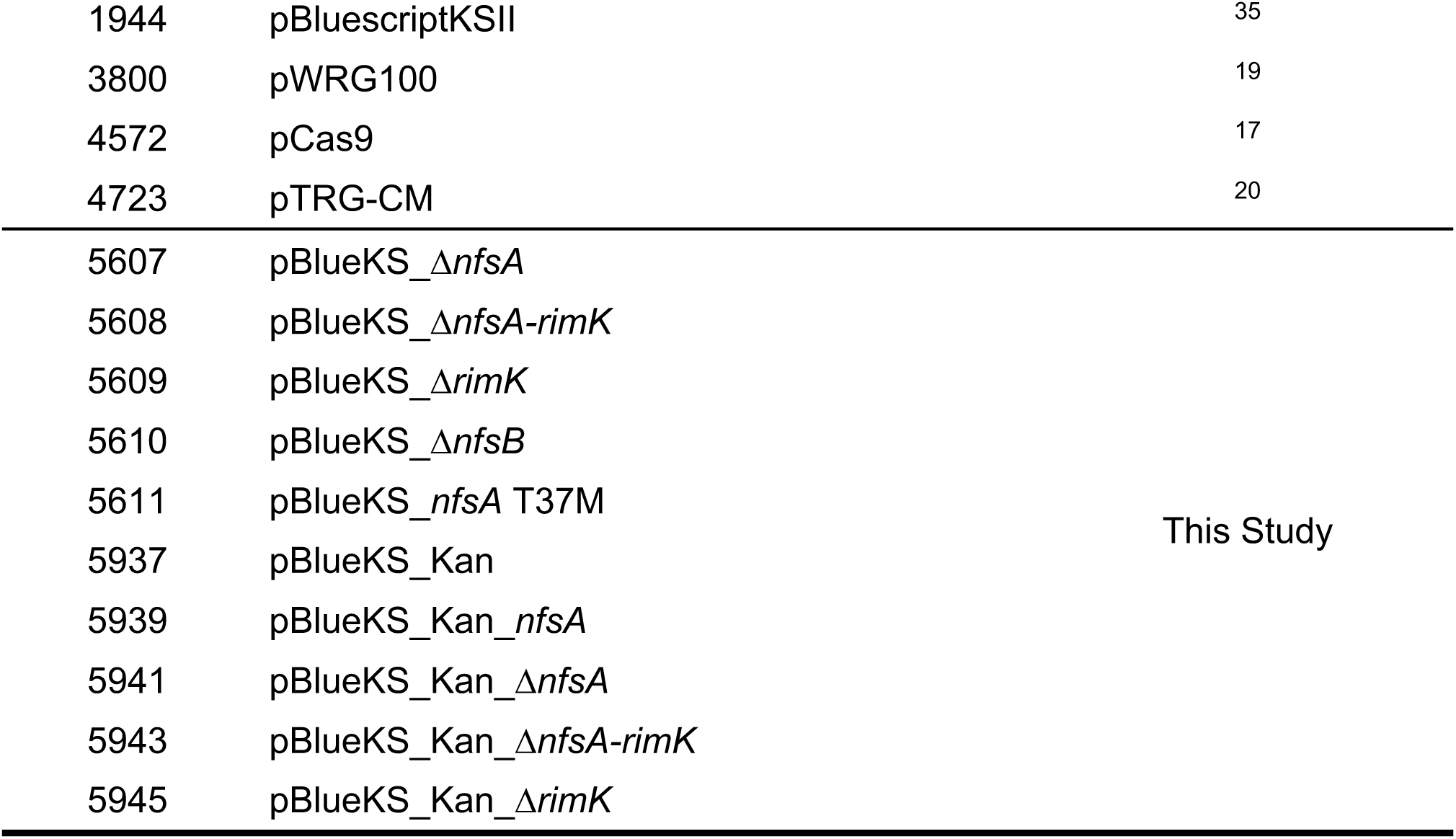
Strains and plasmids used throughout this study

**Table S2.**
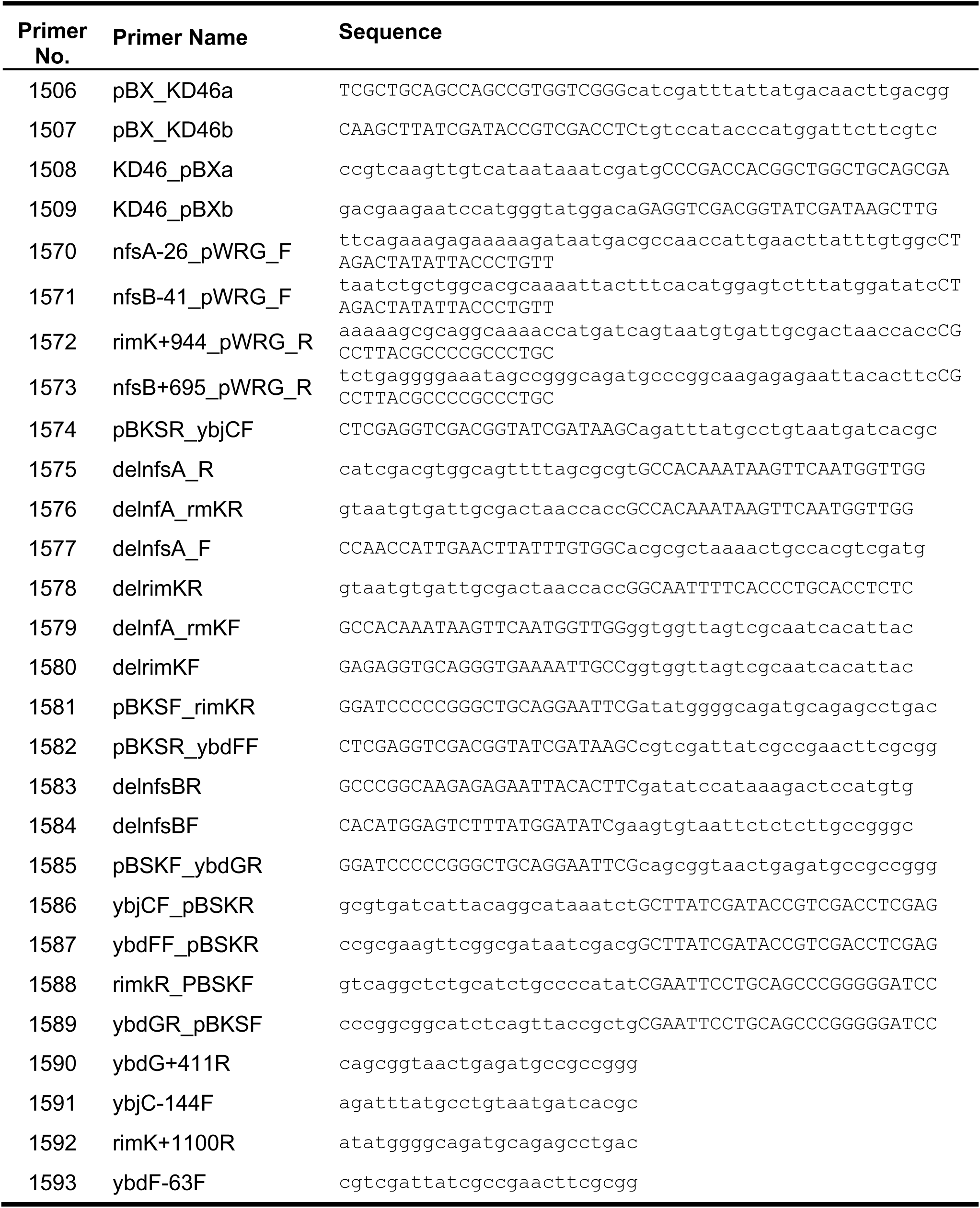
Primers used during this study

**Table S3.**
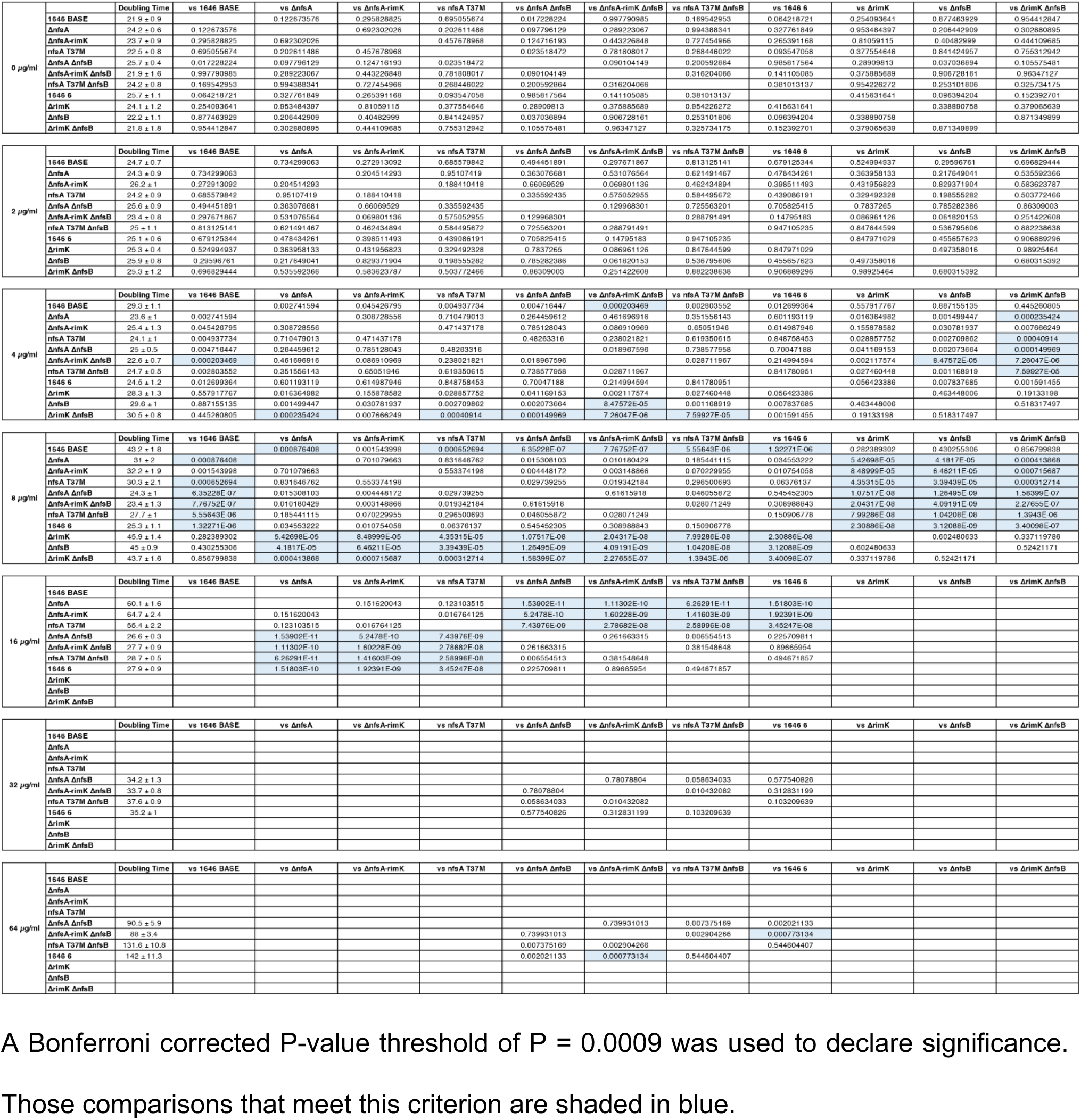
P-value matrix for pairwise comparison of doubling times for all PAT1646 derived strains.

**Table S4.**
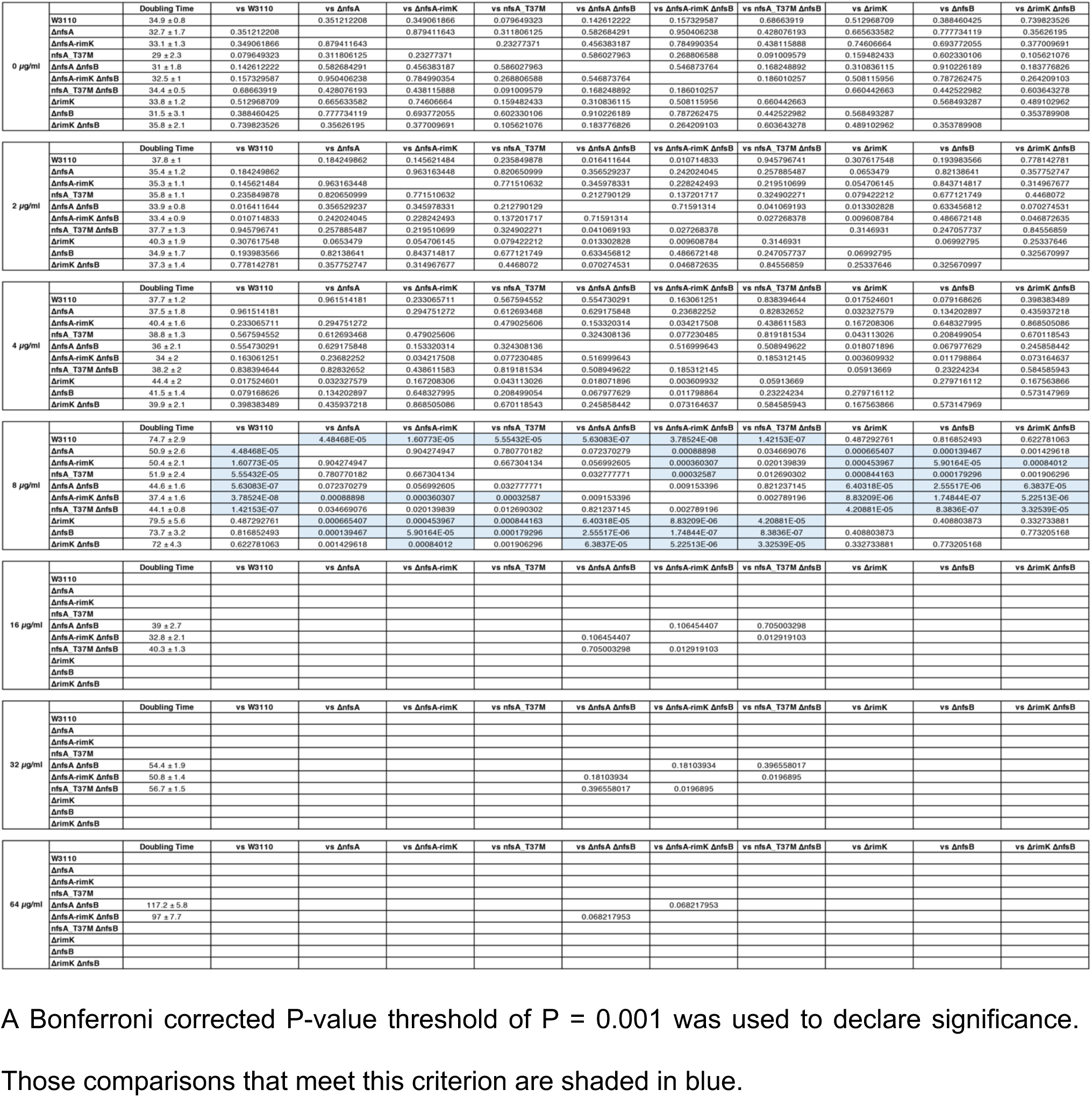
P-value matrix for pairwise comparison of doubling times for all W3110 derived strains for data shown in Table 2.

## Figures and Legends

**Figure S1.**
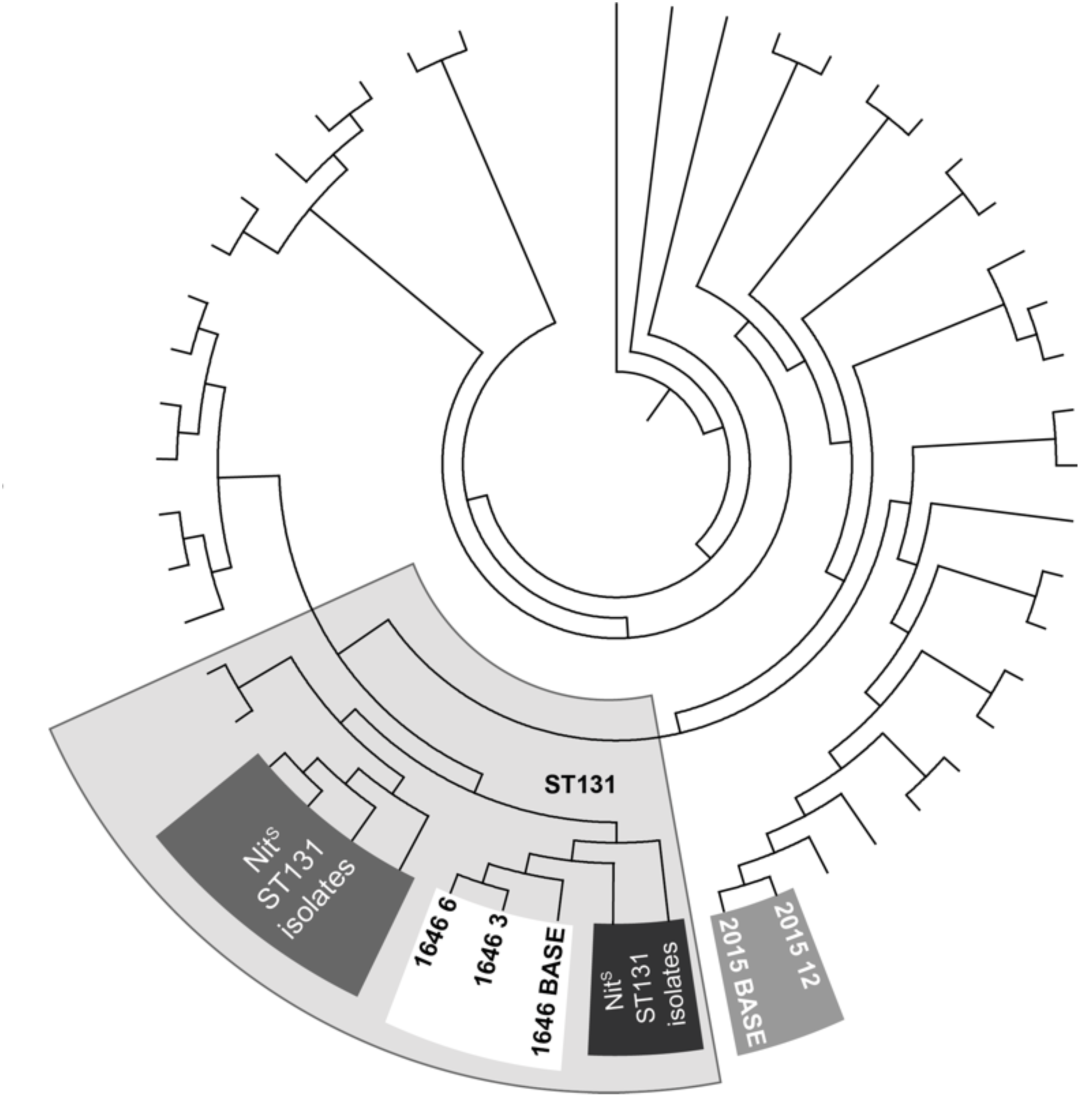
Phylogenetic tree based on the output of cgMLST analysis showing the relatedness of PAT1646 and PAT2015 isolates. The large opaque grey zone shows the number of ST131 isolates identified in Mowbray et al (2022). Two other ST131 isolates were identified but exhibited the Nit^R^ phenotype from the first point of isolation (dark grey box). The method of cgMLST used has been previously described ^15,36^.

**Figure S2.**
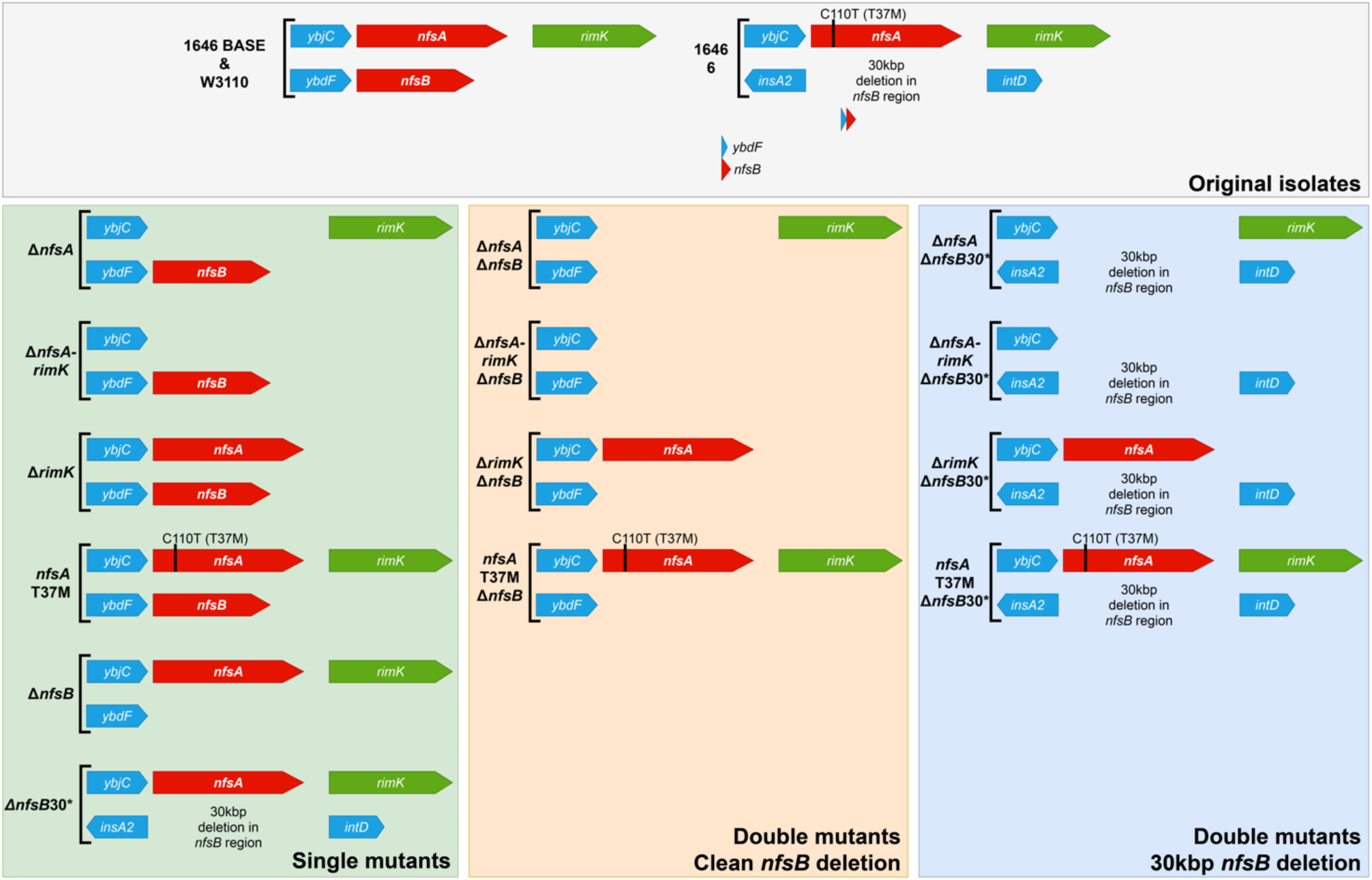
Schematic representation of the genotypes of all mutant combinations made in either 1646 BASE or W3110. Where possible all mutations were made in both genetic backgrounds. The Δ*nfsB30* mutation could not be reconstructed in W3110 due to the lack of synteny upstream of *nfsB* in the parental genomes.

